# COMPARATIVE CHARACTERISTICS OF THE ACCUMULATION OF DIFFERENT VARIANTS OF THE SARS-COV-2 VIRUS (WUHAN, DELTA, OMICRON) IN THE ORGANS OF MODEL ANIMALS

**DOI:** 10.1101/2024.11.12.623177

**Authors:** A.S. Chernov, V.A. Kazakov, I.S. Gogleva, S.V. Savenko, V.N. Schukina, S.Ya. Loginova, S.V. Borisevich, I.V. Smirnov, G.B. Telegin, A.A. Belogurov

## Abstract

**Background:** The genotypic variability of the SARS-CoV-2 virus has proven to be extremely high, and the emergence of new strains raises concerns about their possible high virulence, transmissibility, and ability to bypass responses of the body’s immune system induced by previous infection or vaccination. Therefore, one of the main tasks is to study the pathogenesis of various variants of the virus using experimental animal biomodels of SARS-CoV-2 to quickly find methods and approaches to fighting new viruses.

**Methods:** 60 humanized mice of the C57BL/6-Tgtn (CAG-human AEC2-IRES-Luciferase-WPRE-polyA) line (hACE2) were used. Mice were infected intranasally at different doses with three variants of the SARS-CoV-2 virus: Wuhan, Delta and Omicron.

**Results:** We showed that humanized hACE2 mice, when infected with all three variants of the SARS-CoV-2 virus, showed typical pathological changes in lung consistency comparable to those found in COVID-19 in humans. All mice developed interstitial pneumonia, characterized by inflammatory cell infiltration and thickening of the alveolar septa, characteristic of vascular damage.

**Conclusions:** At a dose of 4 lg plaque-forming unit (PFU), all variants showed 100% mortality. A dose-dependent effect was established only for the Wuhan and Delta variants. In a comparative assessment of different variants of the SARS-CoV-2 virus in a humanized mouse model of hACE2, it was found that the Delta variant leads to more severe damage compared to Wuhan or Omicron.

## INTRODUCTION

Since the SARS-CoV-2 virus first emerged in Wuhan, China, in late 2019^1^, its genetic material has undergone regional changes, some of which have demonstrated the ease of its transmission, the severity of the disease it causes, and the effectiveness of vaccination against it, diagnostic and treatment tools^2,3^. Genomes of the SARS-CoV-2 virus that differ from each other only in detected regions of the genetic sequence are called variants^4^. When variants begin to differ from each other not only genotypically, but also phenotypically, they begin to be called strains^5^. Currently, all variants of the SARS-CoV-2 corona virus are variants.

By mid-summer 2021, the Delta variant was predominant in most countries of the world, including Russia: out of 244 sequenced genomes of the virus, 242 turned out to be the Delta variant. One of the key mutations that give Delta an advantage over other variants is a mutation in the region of the protein where furin cleavage occurs^6^. This mutation increases the proportion of activated viral particles produced after replication by 25% compared to the original variant^7^, and makes it 40–60% more infectious^8^. In terms of contagiousness, it is close to chickenpox.

The Omicron variant was first identified on November 9, 2021 in Botswana and, a few days later, in South Africa^9^. According to the WHO, the new variant spread faster than the previous ones, as evidenced by the data of Russian specialists^10^. This may be explained by the high rate of virus reproduction in the bronchi and nasal epithelium, as well as its ability to bypass the immune system.

Animal biomodels that recapitulate the clinical and pathological features of COVID-19 in humans are an important tool for studying the pathogenesis, mechanisms and transmission routes of the pathogen. Currently, there are several transgenic mouse models in which hACE2 is under the expression of a tissue-specific promoter: K-18-hACE2 mice with the human K-18 promoter^12,13^; HFH4-ACE2 mice with the human HFH4 promoter^14^; mice with the mouse ACE2 promoter [15]; mice AC70, AC22 and AC63 with the CAG promoter^11,16^. Of all the presented models, CAG-hACE2 mice turned out to be the most promising, since the hACE2 protein is expressed not only in the lungs, but also in the brain^17^. Infection in the brain is secondary to the lungs, where pronounced activation of inflammatory mediators is observed already 2 days after infection^17^. For these reasons, CAG-hACE2 mice are highly susceptible to SARS-CoV-2, with the lungs and brain being the main targets.

In connection with the above, the purpose of this study was to compare the pathomorphological features of damage to the lungs of experimental animals when they were infected with different variants of the virus in different doses.

## MATERIALS AND METHODS

### Viruses

Three variants of the SARS-CoV-2 virus were used in the present study:

1. SARS-CoV-2 virus, variant hCoV-19/Russia/GAM-Omicron/2021 (genetic line B.1.1.529, Omicron)
2. SARS-CoV-2 virus, variant hCoV-19/Russia/SPE-RII-32661V-2021 (genetic line B.1.617. (Indian variant Delta) B. 1.617.3 – according to classification dated May 12, 2021. Characteristic mutations GISAID EPI_ISI_1797437).
3. SARS-CoV-2 virus, variant B (lineage B.1.1, Wuhan)

### Cell culture

A permanent culture of African green monkey kidney cells, Vero Cl008, was used in the experiments. For growth and maintenance, Eagle’s medium (MEM) with Hanks’ saline solution containing 7.5% and 2% fetal calf serum, respectively, was used.

### Laboratory animals

We used 60 mice (females and males) 8-10 weeks of age of the CAG-hACE2-IRES-Luc-Tg line (Cat. NO. NM-TG-200002) obtained from the Shanghai Center for Model Organisms^18^. The animals were kept in standard conditions of the Laboratory Animal Nursery of the Institute of Bioorganic Chemistry of the Russian Academy of Sciences (UNU “Bio-model” of the Institute of Bioorganic Chemistry of the Russian Academy of Sciences; Bioresource collection “Collection of laboratory rodents of the SPF category for fundamental, biomedical and pharmacological research”, contract No. 075-15-2021-1067). All experiments and manipulations with animals were approved by the Institutional Committee for the Care and Use of Laboratory Animals (IACUC No.746/20 dated 20/02/21).

### Infection of mice with SARS-CoV-2

Mice were infected with the virus in a specialized laboratory (biosafety level BSL-3) after 3 days of adaptation. Animals were infected intranasally at a dose range of 1–4 lg PFU in a total volume of 20 μl. Infected animals were observed for 12 days, while the animals appearance, behavior, and appetite were monitored. In mice that died during the observation period and were euthanized on day 12, the level of accumulation of the SARS-CoV-2 virus in the internal organs was assessed by the formation of negative colonies in the Vero Cl008 cell culture.

### Titration of the SARS-CoV-2 virus

The infectious activity of the virus was assessed in Vero Cl008 cell culture by the formation of negative colonies under an agar coating^19^. A cell suspension with a density of 200 thousand/ml was added to sterile plastic bottles for cell culture and incubated for 24 hours in a CO_2_ incubator in an atmosphere of 5% CO_2_ at 37°C until a continuous monolayer was formed. Vials with a continuous monolayer were used to determine the infectious titer of the virus. For each tested dose of the drug and control, at least 2 bottles were used. Virus adsorption time was 60 min.

The primary (secondary) agar coating was prepared on Earle’s glucose-salt solution containing l-glutamine, amino acid vitamin complex, antibiotics and fetal calf serum. The incubation time of the infected cell monolayer is 3-5 days, after 3 days the 1st coating, after 24 hours - the 2nd and on the 5th day counting negative colonies (for option B and Delta) and 5-7 days for the Omicron option.

Add 6 ml of neutral red solution (per 100 ml of coating) to the secondary agar coating instead of an equal volume of water.

Calculation of the biological activity (A) of the virus strain culture in PFU/ml was carried out using the formula:

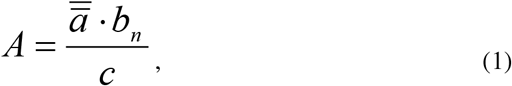

where: =a is the average weighted number of plaques per bottle;

b_n_ – degree of highest dilution;

c – volume of inoculum, ml.

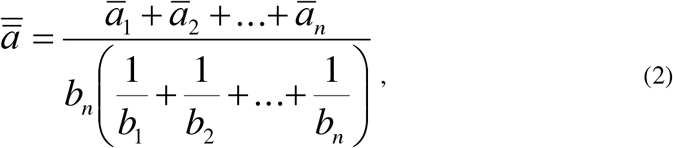

where: -a1….-an – average number of plaques in 1 to n dilution of the test material;

b1…… bn – degree of dilution of the material under study.

### Histology

Lung samples were fixed in a 10% neutral formalin solution for 2 weeks, after which 1 fragment was cut out from each lobe of the lungs. The samples were washed in running water, dehydrated in ethanol of increasing concentration and embedded in paraffin. Paraffin sections 4-5 μm thick, stained with hematoxylin and eosin, were examined using conventional light microscopy on an AxioScope.A1 microscope (Carl Zeiss, Germany). Microphotographs of histological preparations were obtained using a high-resolution camera Axiocam 305 color (Carl Zeiss, Germany) and ZEN 2.6 lite software (Carl Zeiss, Germany).

The severity of various inflammatory phenomena in the lungs was assessed using a semi-quantitative method (in points) according to the following scale: 0 – within normal limits; 1 – weak; 2 – average; 3 – strong.

### Statistical data processing

Statistical processing of the results was carried out using Statistica software (StatSoft®, v.12.6, Tulsa, OK, USA). Results are presented as mean ± SD.

## RESULTS

### Sensitivity of hACE2 mice to different variants of the SARS-CoV-2 virus

The sensitivity of hACE2 mice to the SARS-CoV-2 virus, variant B (Wuhan) was assessed during intranasal infection at doses of 1, 2, 3, 4 lg PFU (Table 1). Monitoring the condition of the animals for 12 days showed that the behavior, mobility and appetite of the infected animals did not differ from the physiological norm. None of the infected animals showed obvious clinical symptoms.

**Table 1.** Assessment of the viral load in the lungs of dead mice and the sensitivity of hACE2 mice to the SARS-CoV-2 virus, variant B (Wuhan) during intranasal infection.

| **Infection dose, lg PFU** | **Frequency of animal deaths** | **Animal death, %** | **Average life time before death, days** | **Level of virus accumulation in the lungs, PFU/g** |
| --- | --- | --- | --- | --- |
| 4 | 5/5 | 100 | 4.0±0 | 2.6±0.3 |
| 3 | 5/5 | 100 | 4.0±0 | 2.9±0.2 |
| 2 | 2/5 | 40 | 9.8±3,9 | 1.3±0.2 |
| 1 | 2/5 | 40 | 11.3±2.8 | 1.5±0.2 |

When mice were intranasally infected with hACE2 at a dose of 3-4 lg PFU, 100% death of mice was recorded. When using the virus at a dose of 1-2 lg PFU, the survival rate was 60%. In all dead mice, the virus was detected in the lung tissues (Table 1). When infected at a dose of 3 lg PFU in mice that died on day 6, the level of viral load was 3.4 PFU/g, while on day 7 it was 2.6 PFU/g. The average rate of virus accumulation in this group was 2.9±0.2 PFU/g (Table 1). When calculating LD50 using the Van der Waerden method, the value was 178 ^x^/:3 PFU/mouse.

When hACE2 mice were intranasally infected with the SARS-CoV-2 virus, variant B (Wuhan), at a dose of 3 lg PFU on the 3rd day after infection (euthanasia), in the brain tissue the level of pathogen accumulation was 5.12 lg PFU/g, in the lungs less 1.0 lg PFU/g (Table 2).

**Table 2.** Variant B, infection dose 3 lg PFU, cervical dislocation.

| **Days after infection** | **Brain** | **Lungs** |
| --- | --- | --- |
| 1 | 0,0±0,0 | 0,0±0,0 |
| 2 | 0,0±0,0 | 0,0±0,0 |
| 3 | 5,12 lg PFU/g | 1 lg PFU/g<br>(the virus was found in 10% of the homogenate suspension) |

In animals that died on the 5th day, the level of virus accumulation in the brain was 6.8 lg PFU/g, and in the lungs only about 2.0 lg PFU/g (Table 3). Thus, the accumulation of the virus in brain tissue is much higher than in the target organ. Viral accumulation was also detected in the liver, spleen and heart (Table 3).

**Table 3.** Accumulation of virus variant B (Wuhan) in different organs of infected mice.

| Organs | PFU/g |
| --- | --- |
| Brain | 6,8±0,03 |
| Lung | 2,01±0,15 |
| Liver | 2,85±0,06 |
| Spleen | 3,10±0,14 |
| Heart | 1,87±0,19 |
| Kidneys | Not identified |

Hematological and biochemical blood parameters of healthy and infected hACE2 mice did not reveal any differences (Table 4).

**Table 4.**
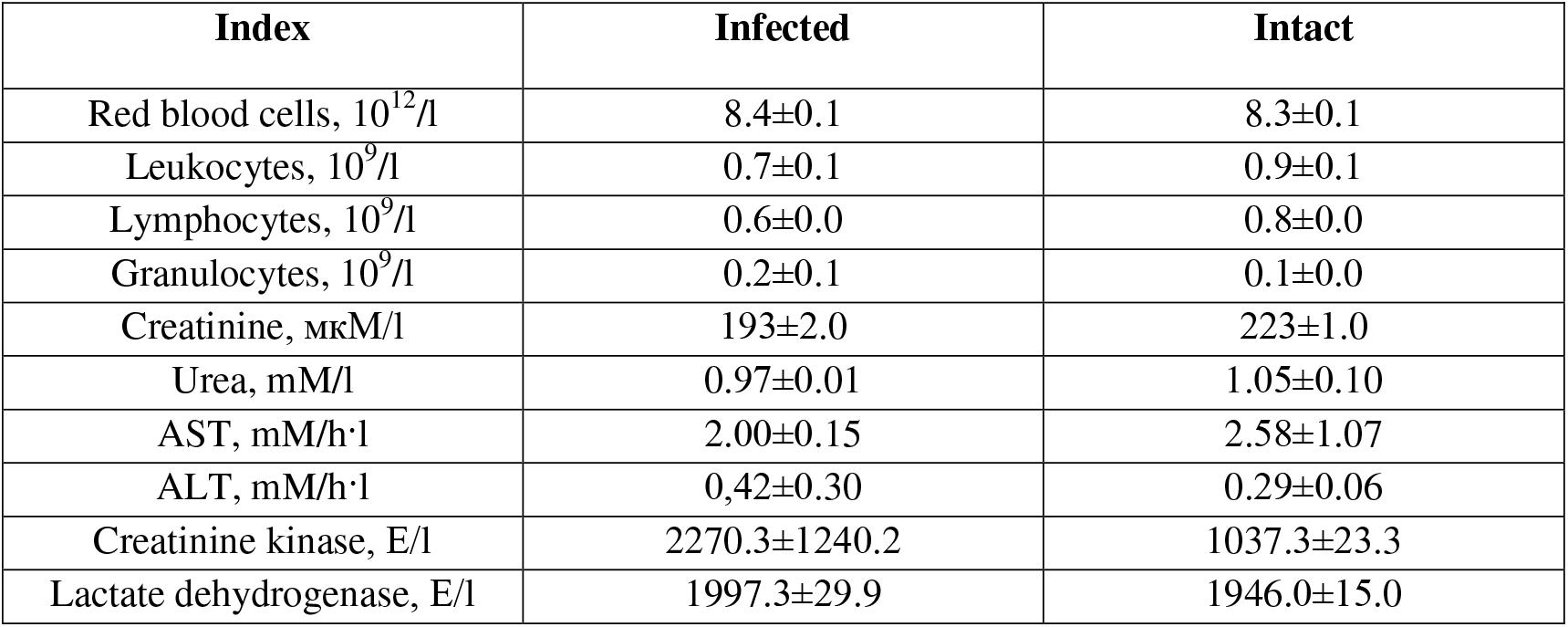
Hematological and biochemical parameters (in living) C57BL/6 hAEC2-TG+ mice on the 3rd day after infection with the SARS-CoV-2 virus, variant B (Wuhan) at a dose of 3 lg PFU.

Studies were conducted to assess the viral load in the lungs of dead hACE2 mice infected intranasally with the SARS-CoV-2 virus, variant hCoV-19/Russia/SPE-RII-32661V-2021 (Delta, line B.1.617.3, Indian variant) (Table. 2). When intranasally infected with the SARS-CoV-2 virus, at a dose of 3 – 4 lg PFU, 100% death of animals was detected, the average life time of mice before death was 5.75 days (Table 5).

**Table 5.** Assessment of the viral load in the lungs of dead mice and the sensitivity of hACE2 mice to the SARS-CoV-2 virus, variant hCoV-19/Russia/SPE-RII-32661V-2021 (Delta, line B.1.617.3, Indian variant) with intranasal infection.

| Infection dose, lg PFU | Frequency of animal deaths | Animal death, % | Average life time before death, days | Level of virus accumulation in the lungs, PFU/g |
| --- | --- | --- | --- | --- |
| 4 | 5/5 | 100 | 5.6±0.2 | 2.0±0.3 |
| 3 | 5/5 | 100 | 5.4±0.2 | 1.7±1.0 |
| 2 | 3/5 | 60 | 5.3±0.3 | 1.3±1.0 |
| 1 | 3/5 | 60 | 6.7±0.3 | 1.3±1.0 |

When animals were infected at a dose of 1-2 lg PFU, death was at the level of 60%, the average life time of mice before death was 6.7 days and 5.3 days, respectively. When mice were infected at a dose of 4 lg PFU, 40% had an accumulation level in the lungs of 1.3 PFU/g, and 60% had an accumulation level of 2.6 PFU/g. The average accumulation of virus in the lungs of animals in this group was 2.0±0.3 PFU/g.

When mice were infected at a dose of 3 lg PFU, the level of accumulation in the lungs was 1.3 PFU/g in 60%, and 2.1 PFU/g in 40%. The average accumulation of virus in the lungs of animals in this group was 1.7±0.6 PFU/g.

When mice were infected at a dose of 1–2 PFU, 100% had an accumulation level in the lungs of 1.3 PFU/g. When calculating LD50 using the Van der Waerden method, the value was 79.56 ^x^/:2.7 PFU/mouse, 1.9±0.4 PFU.

Intranasal infection of mice with the SARS-CoV-2 virus, variant hCoV-19/Russia/GAM-Omicron/2021 (line B.1.1.529, Omicron) was carried out at a dose of 1 - 4 lg PFU in 20 μl per mouse. Monitoring the condition of the animals showed that the behavior, mobility and appetite of the infected animals did not differ from the physiological norm.

During intranasal infection of hACE2 mice with the SARS-CoV-2 virus, variant hCoV-19/Russia/GAM-Omicron/2021 (line B.1.1.529) at a dose of 4 lg PFU, 100% death of animals was detected, the average life time of mice before death was 7.6 days (Table 6).

**Table 6.** Assessment of the sensitivity of hACE2 mice to the SARS-CoV-2 virus, variant hCoV-19/Russia/GAM-Omicron/2021 (line B.1.1.529) during intranasal infection

| Infection dose, lg PFU | Frequency of animal deaths | Animal death, % | Average life time before death, days | Level of virus accumulation in the lungs, PFU/g |
| --- | --- | --- | --- | --- |
| 4 | 5/5 | 100 | 7.6±0.2 | 3.3±0.1 |
| 3 | 3/5 | 60 | 7.6±0.3 | 3.3±0.1 |
| 2 | 4/5 | 80 | 9.3±0.3 | 2.9±0.1 |
| 1 | 3/5 | 60 | 9.3±0.9 | 2.3±0.1 |

When infected at a dose of 1 and 3 lg PFU, 60% of the animals died, the average life time of mice before death was 7.6 days and 9.3 days, respectively. When mice were infected at a dose of 2 PFU, the death rate was 80%; the average life time of mice before death was 9.3 days.

It should be noted that at an infection dose of 1-2 lg PFU, animals that died on days 9, 10 and 11 after infection with the SARS-CoV-2 virus over the last three days were inactive, with heavy breathing and wheezing.

Studies were conducted to assess the viral load in the lungs of dead hACE2 mice infected intranasally with the SARS-CoV-2 virus, variant hCoV-19/Russia/GAM-Omicron/2021 (line B.1.1.529) (Table 6). From the data presented in Table 3, it follows that the level of virus accumulation in the target organ is practically independent of the dose of infection. When calculating LD50 using the Van der Waerden method, the value was 125.9 ^x^/:3.7 PFU/mouse, 2.1±0.6 lg PFU.

### Histology results

Unlike healthy mice, in the lungs of hACE2 mice infected with the Wuhan variant of SARS-CoV-2 at doses of 2-4 lg PFU and dying on days 6-7, the following pathomorphological picture was noted: diffuse pronounced congestion of large vessels, as well as blood vessels microvasculature with symptoms of stasis and sludge of erythrocytes (Fig. 1, black arrows). In the walls of the alveoli, the focal presence of a few macrophages, neutrophils and lymphocytes was noted, and focal perivascular mononuclear infiltration was found around individual vessels (Fig. 1, yellow arrow). In the lumen of individual alveoli in different lobes of the organ, hyaline-like membranes and transudate were found. In the lumen of most bronchi, noticeable was the pronounced desquamation of the respiratory epithelium with the initial destruction of the integrity of the bronchial wall. The pathomorphological changes described above in autopsy specimens of the lungs of hACE2 mice infected with the Wuhan strain of SARS-CoV-2 at doses of 2-4 lg PFU are clearly dose-dependent.

**Figure 1.**
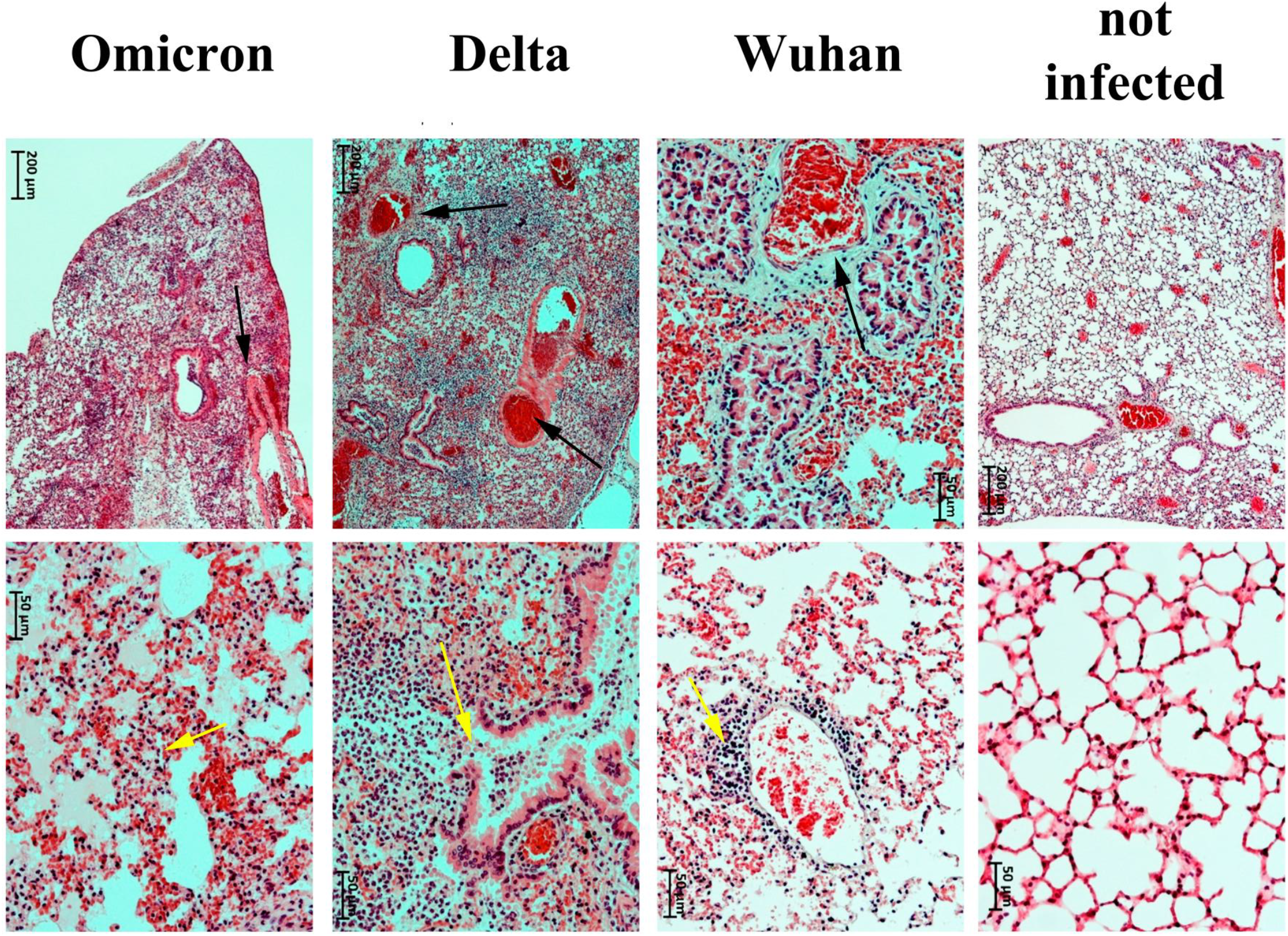
Lung fragments from hACE2 mice intact and infected with the Wuhan variant of SARS-CoV-2, Delta and Omicron, which died within 5-8 days after infection. Staining with hematoxylin and eosin (magnification 50x and 200x).

Half of the hACE2 mice infected with the Delta variant at doses of 2-4 lg PFU and dying after 5-6 days, as a rule, had numerous large foci of necrosis in different lobes of the lungs as a result of perforation of the bronchial wall (Fig. 1, yellow arrow). Foci of necrosis were abundantly infiltrated with segmented neutrophils; numerous mononuclear cells were also found. In foci of necrosis, the interalveolar septa are destroyed; hyaline-like membranes and transudate can be found in the lumen of intact alveoli. In cases where necrosis was not observed, the picture was generally similar to that noted when mice were infected with the Wuhan variant - pronounced disturbances at the level of the microvasculature with phenomena of stasis and sludge of erythrocytes, hyaline-like membranes in the lumen of the alveoli, massive areas of transudates. Insignificant or moderate mononuclear infiltration was found in the walls of the alveoli, and focal mononuclear infiltrates were found in the perivascular space. In general, Delta is characterized by a dose dependence in terms of the volume of damage to lung tissue and the severity of the pathomorphological picture.

In the lungs of hACE2 mice infected with the Omicron strain at doses of 1-4 lg PFU and dying after 7-9 days, a generally similar pattern of pathomorphological changes was observed as when infected with the Wuhan strain of SARS-CoV-2 at a dose of 3 lg PFU without obvious dose dependence. In all cases, there were pronounced disturbances at the level of the microvasculature with phenomena of stasis and sludge of erythrocytes (Fig. 1, black arrow). In the lumen of the alveoli, hyaline-like membranes and massive areas of transudates were found. Moderate or pronounced mononuclear infiltration was noticeable in the walls of the alveoli; focal mononuclear (neutrophil - in a single case) infiltrates were found in the perivascular space. In 1 male rat infected with the Omicron strain at a dose of 1 lg PFU, extensive foci of necrosis were observed in 2 lobes of the lungs with destruction of the interalveolar septa and abundant infiltration of tissues with segmented leukocytes. In both cases, bronchial perforation occurred in the foci of necrosis. Considering what was described above, no obvious dose-dependence in doses of the Omicron strain 1-3 lg PFU was detected. At a high dose of infection (4 lg PFU), all animals with a typical pathomorphological picture in the lungs died.

Surviving hACE2 mice infected with different variants of SARS-CoV-2 and euthanized on day 12 of the study did not show any obvious pathological changes in the lungs.

Summary data on the frequency of occurrence of pathohistological signs in the lungs of hACE2 mice infected with the Wuhan strain of SARS-CoV-2 at a dose of 1-4 lg PFU, Omicron and Delta strains at doses of 1-4 lg PFU are presented in Table. 7.

**Table 7.**
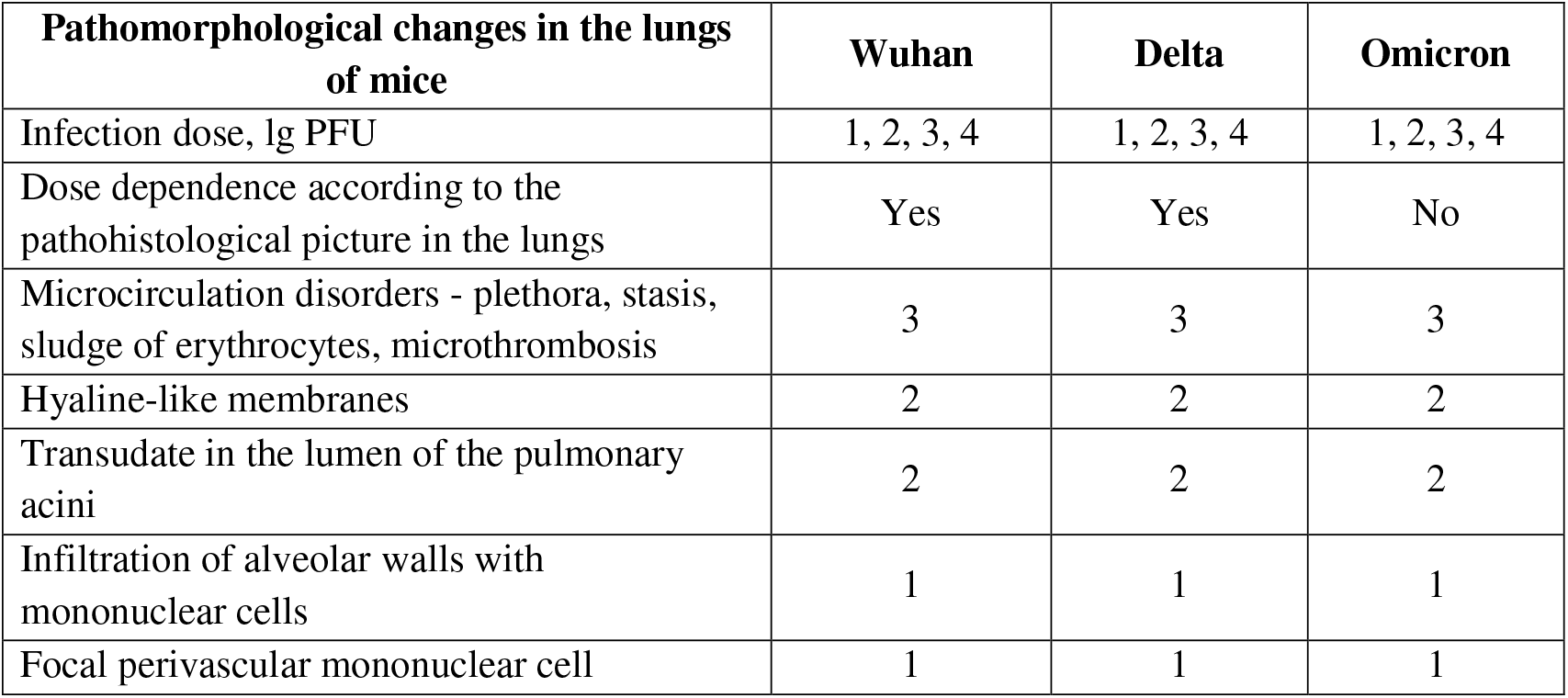
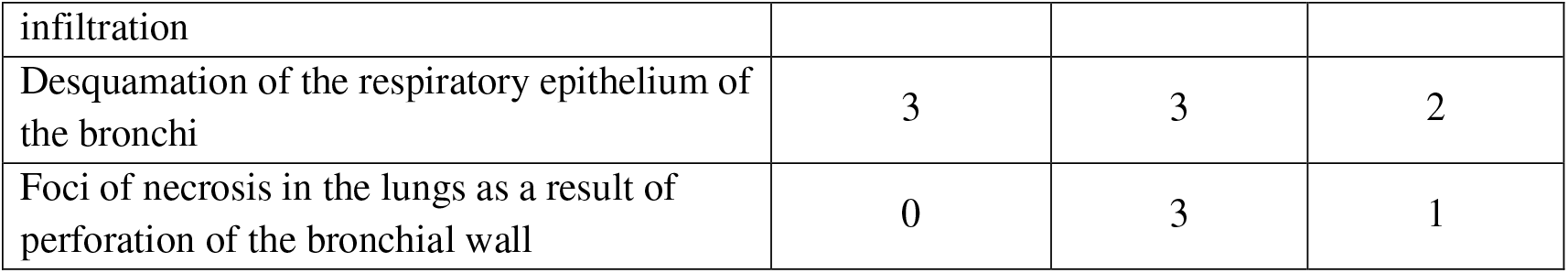
Frequency of occurrence of pathomorphological changes (in points) in the lungs of hACE2 mice infected with various variants of SARS-CoV-2, which died within 5-9 days after infection.

## DISCUSSION

As it became known, linear mice are not susceptible to the original variant of SARS-CoV-2 that caused COVID-19. The key receptor for interaction with SARS-CoV-2 in humans is ACE2, interaction with which occurs through the receptor binding domain (RBD) of the virus S protein. Murine ACE2 (mACE2) prevents infection by SARS-CoV-2, which became the basis for the use of a special humanized transgenic mouse line C57BL/6 hAEC2-TG+ expressing human ACE2^20^. Our virological and pathological studies showed that all virus variants cause productive infection in hACE2 mice, which coincides with previous studies^18,20,21^. The use of the virus in doses of 3-4 lg PFU led to 100% lethal outcome of all infected hACE2 mice, which indicates the high sensitivity and adequacy of the model used. These data make this model the most preferable for research in the field of SARS-CoV-2, in contrast to the same Syrian hamsters, for which high doses of the virus (5-6 lg PFU) are used, but at the same time they develop only moderate signs of clinical diseases and pathologies of the lungs^17,22,23,24^.

A histological study of autopsy specimens of the lungs of hACE2 mice infected with different variants of SARS-CoV-2 demonstrated a typical pathomorphological picture in the target organ, characterized by pronounced diffuse congestion of the microvasculature with phenomena of stasis and sludge of erythrocytes, signs of thrombus formation, as well as infiltration of alveolar walls by inflammatory cells, the presence of hyaline-like membranes and transudate in the lumen of the pulmonary acini. In light of the 50% probability of perforation of the bronchial walls, the most aggressive variant of the SARS-CoV-2 virus during intranasal infection of hACE2 mice appears to be the Delta variant, which, along with the Wuhan variant, showed a dose-dependent nature of target organ damage. As was established for the Wuhan variant, the accumulation of the virus occurs not only in the target organ, but also in the brain tissue, and to a much greater extent. The results obtained correlate well with studies in humans^25,26,27,28^, and, as is known, in people who had COVID-19 of increased risk of long-term neurologic disorders^29,30,31^.

In our study, no hematological and biochemical changes in blood parameters were detected. However, some studies suggest to the panel of hematological and biochemical markers that can predict and assess the risk of disease severity and mortality in patients with COVID-19^32,33^.

It is possible that for the presented animal model it is necessary to use a different method of infection, since during intranasal infection the virus travels the shortest route along the nerve to the brain. And with such a level of virus in the brain, it will be difficult to stop the virus from reproducing using therapeutic agents.

## CONCLUSION

Mice C57BL/6 hAEC2-TG+ (C57BL/6-Tgtn(CAG-human AEC2-IRES-Luciferase-WPRE-polyA) during intranasal infection are highly susceptible to the SARS-CoV-2 virus, variant B (Wuhan), hCoV-19 variant/Russia/SPE-RII-32661V-2021 (delta, line B.1.617.3, Indian variant), variant hCoV-19/Russia/GAM-Omicron/2021 (line B.1.1.529).

In a comparative assessment of various variants of the SARS-CoV-2 virus on the humanized mouse model C57BL/6 hAEC2-TG+, it was found that the Delta variant leads to more severe damage compared to Wuhan or Omicron, which correlates well with clinical observations in humans and the results of the study [23]. We conducted a pathohistological study of the lungs of infected mice. Omicron is close to the Delta variant in terms of the severity of pathomorphological changes in the lungs, but does not have obvious dose-dependent characteristics of a damaging effect on the target organ, unlike the Wuhan and Delta variants.

The humanized C57BL/6 hAEC2-TG+ mice used in this work adequately model the pathological processes of SARS-CoV-2 in humans, and can be successfully used in testing diagnostic tools and therapy for infectious diseases.

## ETHICS APPROVAL

All experiments and manipulations with animals were approved by the Institute Committee for the Care and Use of Laboratory Animals of the Institute of Bioorganic Chemistry of the Russian Academy of Sciences (IACUC No. 746/20 dated 02/20/21).

## INFORMED CONSENT

Not applicable due to the design of the study.

## AVAILABILITY OF DATA AND MATERIALS

The datasets generated and/or analyzed during the current study are not publicly available but are available from the corresponding author upon reasonable request.

## AUTHORS’ CONTRIBUTIONS

Conceptualization: Aleksandr Chernov, Georgii Telegin, Alexey Belogurov; methodology: Aleksandr Chernov, Vitaly Kazakov, Irina Gogleva, Sergei Savenko, Svetlana Loginova, Veronika Schukina; formal analysis: Georgii Telegin, Alexey Belogurov; investigation: Aleksandr Chernov, Alexey Minakov, Irina Gogleva, Sergei Savenko, Veronika Schukina; data curation, Aleksandr Chernov, Vitaly Kazakov, Ivan Smirnov; writing-original draft preparation, Aleksandr Chernov, Svetlana Loginova, Alexey Belogurov; writing-review and editing, Georgii Telegin, Alexey Belogurov; supervision, Aleksandr Chernov. All authors have read and agreed to the published version of the manuscript.

## REFERENCES

1. Zhu H, Wei L, Niu P. The novel coronavirus outbreak in Wuhan, China. Glob Health Res Policy 2020; 5:6.

2. Oberemok VV, Laikova KV, Yurchenko KA, Fomochkina II, Kubyshkin AV. SARS-CoV-2 will continue to circulate in the human population: an opinion from the point of view of the virus-host relationship. Inflamm Res 2020; 69(7):635–640.

3. Saberiyan M, Karimi E, Khademi Z, Movahhed P, Safi A, Mehri-Ghahfarrokhi A. SARS-CoV-2: phenotype, genotype, and characterization of different variants. Cell Mol Biol Lett 2022; 17;27(1):50.

4. Aleem A, Akbar Samad AB, Vaqar S. Emerging Variants of SARS-CoV-2 And Novel Therapeutics Against Coronavirus (COVID-19). In: StatPearls [Internet]. Treasure Island (FL): StatPearls Publishing 2023; 34033342.

5. Dos Santos WG. Impact of virus genetic variability and host immunity for the success of COVID-19 vaccines. Biomed Pharmacother 2021; 136:111272.

6. Liu Y, Liu J, Johnson BA, Xia H, Ku Z, Schindewolf C, Widen SG, An Z, Weaver SC, Menachery VD, Xie X, Shi PY. Delta spike P681R mutation enhances SARS-CoV-2 fitness over Alpha variant. bioRxiv [Preprint]. 2021; 5:2021.08.12.456173.

7. Saito A, Irie T, Suzuki R. et al. Enhanced fusogenicity and pathogenicity of SARS-CoV-2 Delta P681R mutation. Nature 2022; 602, 300–306.

8. Biswas B, Chattopadhyay S, Hazra, S. et al. COVID-19 pandemic: the delta variant, T-cell responses, and the efficacy of developing vaccines. Inflamm Res 2022; 71, 377–396.

9. Mohapatra RK, Sarangi AK, Kandi V, Azam M, Tiwari R, Dhama K. Omicron (B.1.1.529 variant of SARS-CoV-2); an emerging threat: Current global scenario. J Med Virol 2022; 94(5):1780–1783.

10. Vechorko V., Averkov O., Zimin A. New SARS-CoV-2 Omicron variant — clinical picture, treatment, prevention (literature review). Cardiovascular Therapy and Prevention 2022; 21. 3228.

11. Tseng CT, Huang C, Newman P, et al. Severe acute respiratory syndrome coronavirus infection of mice transgenic for the human Angiotensin-converting enzyme 2 virus receptor. J Virol 2007; 81(3):1162–1173.

12. McCray PB, J., Pewe L, Wohlford-Lenane C, Hickey M, Manzel L, Shi L, et al. Lethal infection of K18-hACE2 mice infected with severe acute respiratory syndrome coronavirus. J Virol 2007; 81(2):813–821.

13. Netland J, Meyerholz DK, Moore S, Cassell M, Perlman S. Severe acute respiratory syndrome coronavirus infection causes neuronal death in the absence of encephalitis in mice transgenic for human ACE2. J Virol 2008; 82(15):7264–7275.

14. Menachery VD, Yount BL, J., Sims AC, Debbink K, Agnihothram SS, Gralinski LE, et al. SARS-like WIV1-CoV poised for human emergence. Proc Natl Acad Sci USA 2016; 113(11):3048–3053.

15. Yang XH, Deng W, Tong Z, Liu YX, Zhang LF, Zhu H, et al. Mice transgenic for human angiotensin-converting enzyme 2 provide a model for SARS coronavirus infection. Comp Med 2007; 57(5):450–459.

16. Yoshikawa N, Yoshikawa T, Hill T, Huang C, Watts DM, Makino S, et al. Differential virological and immunological outcome of severe acute respiratory syndrome coronavirus infection in susceptible and resistant transgenic mice expressing human angiotensin-converting enzyme 2. J Virol 2009; 83(11):5451–5465.

17. Asaka MN, Utsumi D, Kamada H, Nagata S, Nakach, Y, Yamaguchi T, Kawaoka Y, Kuba K, Yasutom, Y. Highly susceptible SARS-CoV-2 model in CAG promoter-driven hACE2-transgenic mice. JCI Insight 2021; 6, e152529.

18. Tretiakova DS, Alekseeva AS, Onishchenko NR, Boldyrev IA, Egorova NS, Vasina DV, Gushchin VA, Chernov AS, Telegin GB, Kazakov VA, Plokhikh KS, Konovalova MV, Svirshchevskaya EV, Vodovozova EL. Proof-of-Concept Study of Liposomes with a Set of SARS-CoV-2 Viral Peptidic T-Cell Epitopes as a Vaccine. Russ J Bioorg Chem 2022; 48(Suppl 1):S23–37.

19. Loginova SYa, Shchukina VN, Savenko SV, Borisevich SV. Antiviral activity of Kagocel® in vitro against virus SARS-CoV-2. Antibiot Chemother 2020; 65:3–4.

20. Chernov AS, Minakov AA, Kazakov VA, Rodionov MV, Rybalkin IN, Vlasik TN, Yashin DV, Saschenko LP, Kudriaeva AA, Belogurov AA, Smirnov IV, Loginova SY, Schukina VN, Savenko SV, Borisevich SV, Zykov KA, Gabibov AG, Telegin GB. A new mouse unilateral model of diffuse alveolar damage of the lung. Inflamm Res 2022; 17:1–13.

21. Chernov AS, Rodionov MV, Kazakov VA, Ivanova KA, Meshcheryakov FA, Kudriaeva AA, Gabibov AG, Telegin GB, Belogurov AA Jr. CCR5/CXCR3 antagonist TAK-779 prevents diffuse alveolar damage of the lung in the murine model of the acute respiratory distress syndrome. Front Pharmacol 2024; 21;15:1351655.

22. Francis ME, Goncin U, Kroeker A, Swan C, Ralph R, Lu Y, Etzioni AL, Falzarano D, Gerdts V, Machtaler S, Kindrachuk J, Kelvin AA. SARS-CoV-2 infection in the Syrian hamster model causes inflammation as well as type I interferon dysregulation in both respiratory and non-respiratory tissues including the heart and kidney. PLoS Pathog 2021; 17(7):e1009705.

23. Handley A, Ryan KA, Davies ER, Bewley KR, Carnell OT, Challis A, Coombes NS, Fotheringham SA, Gooch KE, Charlton M, Harris DJ, Kennard C, Ngabo D, Weldon TM, Salguero FJ, Funnell SGP, Hall Y. SARS-CoV-2 Disease Severity in the Golden Syrian Hamster Model of Infection Is Related to the Volume of Intranasal Inoculum. Viruses 2023; 14;15(3):748.

24. Varea-Jiménez E, Aznar Cano E, Vega-Piris L, Martínez Sánchez EV, Mazagatos C, García San Miguel Rodríguez-Alarcón L, Casas I, Sierra Moros MJ, Iglesias-Caballero M, Vazquez-Morón S, Larrauri A, Monge S Comparative severity of COVID-19 cases caused by Alpha, Delta or Omicron SARS-CoV-2 variants and its association with vaccination]. Enferm Infecc Microbiol Clin 2022; 42(4):187–194.

25. Stein SR, Ramelli SC, Grazioli A. et al. SARS-CoV-2 infection and persistence in the human body and brain at autopsy. Nature 2022; 612, 758–763.

26. Poloni TE, Moretti M, Medici V, Turturici E, Belli G, Cavriani E, Visonà SD, Ross, M, Fantini V, Ferrari RR. et al. COVID-19 Pathology in the Lung, Kidney, Heart and Brain: The Different Roles of T-Cells, Macrophages, and Microthrombosis. Cells 2022; 11, 3124.

27. Crunfli F, Carregari VC, Veras FP, Silva LS, Nogueira MH, Antunes ASLM, Vendramini PH, Valença AGF, Brandão-Teles C, Zuccoli GDS, Reis-de-Oliveira G, Silva-Costa LC, Saia-Cereda VM, Smith BJ, Codo AC, de Souza GF, Muraro SP, Parise PL, Toledo-Teixeira DA, Santos de Castro ÍM, Melo BM, Almeida GM, Firmino EMS, Paiva IM, Silva BMS, Guimarães RM, Mendes ND, Ludwig RL, Ruiz GP, Knittel TL, Davanzo GG, Gerhardt JA, Rodrigues PB, Forato J, Amorim MR, Brunetti NS, Martini MC, Benatti MN, Batah SS, Siyuan L, João RB, Aventurato ÍK, Rabelo de Brito M, Mendes MJ, da Costa BA, Alvim MKM, da Silva Júnior JR, Damião LL, de Sousa IMP, da Rocha ED, Gonçalves SM, Lopes da Silva LH, Bettini V, Campos BM, Ludwig G, Tavares LA, Pontelli MC, Viana RMM, Martins RB, Vieira AS, Alves-Filho JC, Arruda E, Podolsky-Gondim GG, Santos MV, Neder L, Damasio A, Rehen S, Vinolo MAR, Munhoz CD, Louzada-Junior P, Oliveira RD, Cunha FQ, Nakaya HI, Mauad T, Duarte-Neto AN, Ferraz da Silva LF, Dolhnikoff M, Saldiva PHN, Farias AS, Cendes F, Moraes-Vieira PMM, Fabro AT, Sebollela A, Proença-Modena JL, Yasuda CL, Mori MA, Cunha TM, Martins-de-Souza D. Morphological, cellular, and molecular basis of brain infection in COVID-19 patients. Proc Natl Acad Sci USA 2022; 119(35):e2200960119.

28. Succi G, Pedrycz W, Bogachuk AP, Tormasov AG, Belogurov AA, Spallone A. The Fallout of Catastrophic Technogenic Emissions of Toxic Gases Can Negatively Affect Covid-19 Clinical Course. Acta Naturae 2022; 14(4):101–110.

29. Xu E, Xie Y, Al-Aly Z. Long-term neurologic outcomes of COVID-19. Nat Med 2022; 28, 2406–2415.

30. Osailan AM. Batarfa M, Aldosari Z, Alghamdi A, Alqahtani F, Alabdullah M, Elnaggar RK Association between COVID-19 exposure and autonomic nervous system dysfunction in apparently healthy adults: an observational study. Eur Rev Med Pharmacol Sci 2024; 28:3420–3429.

31. Monaco S, Massari MG, Renzi A, DI Trani M. COVID-19 post-traumatic stress disorder: the role of ACEs, alexithymia, and attachment in the Italian population. Eur Rev Med Pharmacol Sci 2024; 28:2615–2624.

32. Alizad G, Ayatollahi AA, Shariati Samani A, Samadizadeh S, Aghcheli B, Rajabi A, Nakstad B, Tahamtan A. Hematological and Biochemical Laboratory Parameters in COVID-19 Patients: A Retrospective Modeling Study of Severity and Mortality Predictors. Biomed Res Int 2023; 18:7753631.

33. Pertiwi D, Nisa M, Aulia AP Rahayu,. Hematological and Biochemical Parameters at Admission as Predictors for Mortality in Patients with Moderate to Severe COVID-19. Ethiop J Health Sci 2023; 33(2):193–202.

